# Experimental method for creating skin with acquired appendage dysfunction

**DOI:** 10.1101/2024.09.02.610845

**Authors:** Yuta Moriwaki, Makoto Shiraishi, Qi Shen, Zening Du, Mutsumi Okazaki, Masakazu Kurita

## Abstract

Mammalian skin appendages, such as hair follicles and sweat glands, are essential for both esthetic and functional purposes. Conditions such as burns and ulcers can lead to dysfunction or loss of skin appendages and result in hair loss and dry skin, posing challenges in their regeneration. Existing animal models are insufficient for studying acquired dysfunction of skin appendages without underlying genetic causes. This study aimed to develop more clinically relevant mouse models by evaluating two approaches: keratinocyte transplantation and grafting of skin at varying thicknesses. GFP-expressing keratinocytes were transplanted into ulcers on nude mice, leading to re-epithelialization with minimal skin appendages at 4 weeks after transplantation. However, the re-epithelialized area was largely derived from recipient cells, with the grafted cells contributing to only 1.31% of the area. In the skin grafting model, donor skin from GFP transgenic mice was grafted onto nude mice at three thicknesses: full thickness, 10/1000 inch, and 5/1000 inch. The grafted area of the 5/1000-inch grafts remained stable at 89.5% of its original size 5 weeks after transplantation, ensuring a sufficiently large skin area. The 5/1000-inch grafts resulted in a significant reduction in skin appendages, with a mean of only 3.73 hair follicles per 5 mm, compared with 69.7 in the control group. The 5/1000-inch skin grafting in orthotopic autologous transplantation also showed the achievement of skin surfaces with a minimal number of skin appendages. Therefore, a mouse model with skin grafting demonstrated stability in producing large areas of skin with minimal appendages. In conclusion, these two models with acquired skin appendage dysfunction and no underlying genetic causes provide valuable tools for researching skin appendage regeneration, offering insights into potential therapeutic strategies for conditions involving skin appendage loss.

## Introduction

Mammalian skin appendages are complex mini-organs formed during skin development, which include hair follicles and sweat glands.^1, 2^ Lack or dysfunction of skin appendages can occur due to intrinsic factors such as hormonal influences (androgenic alopecia) and immune responses (alopecia areata), but also external factors such as scars resulting from burns and ulcer healing, as well as skin grafts.^3, 4^ Such conditions cause a number of esthetic and functional complaints, including hair loss and dry skin.^5^

The research and development of new treatments, particularly those aimed at regenerating skin appendages by using local resident cells, require a skin model with declined function and/or a reduced number of skin appendages not caused by genetic or intrinsic factors. However, existing animal models, such as those using sites that naturally lack hair^6–8^ or animals with genetic factors that impede hair growth,^9–11^ do not meet this criterion.

To develop a more clinically relevant model of acquired skin appendage dysfunction, characterized by a quantitative reduction of skin appendages despite the local cells not having any inherent abnormalities, we examined the condition of skin after the formation of a simple ulcer, after the transplantation of cultured keratinocytes onto the ulcer surface, and after grafting of skin with varying thicknesses. For the transplantation procedures, allogeneic transplantation of tissues harvested from mice expressing green fluorescent protein (GFP) in all cells throughout their bodies to immunodeficient nude mice was used to identify the origin of the cells.

## Methods

### Isolation and culture of mouse keratinocytes

Mouse keratinocytes were isolated from the back skin of 3–5-week-old GFP transgenic C57BL/6-Tg (CAG-EGFP) mice (Japan SLC, Inc., Hamamatsu, Japan). The skin specimens were washed three times in phosphate-buffered saline (PBS) and incubated with 0.25% trypsin in PBS for 16–24 h at 4 °C. The epidermis was separated from the dermis using forceps. Keratinocyte culture was started at 37 °C with 5% CO_2_, and maintained on mitomycin C-treated 3T3-J2 feeder cells (a generous gift from the late Dr. Howard Green) in F medium [3:1 (v/v) Ham’s F12 and D-MEM; FUJIFILM Wako Pure Chemical Corporation, Osaka, Japan] supplemented with 5% FBS, 0.4 μg/ml hydrocortisone (Sigma-Aldrich, St. Louis, MO, USA), 5 μg/ml insulin (Sigma), 10 ng/ml EGF (Sigma), 24 μg/ml adenine (Sigma), 8.4 ng/ml cholera toxin (Wako), 100 U/ml penicillin (Wako), 100 μg/ml streptomycin (Wako), and 10 μM Rho-kinase inhibitor Y27632 (Selleck Chemicals, Houston, TX, USA).^12^ The cells were transplanted at passage 2. Imaging was performed using an Olympus IX73 Inverted LED Fluorescence Microscope (Olympus Corporation, Tokyo, Japan).

### Re-epithelialization after ulcer formation

Four-week-old C57BL/6J Jcl (B6) female mice (Nippon Bio-Supp. Center, Tokyo, Japan) were used as recipient animals. Under inhalation anesthesia, a 10 mm circular piece of skin was excised from the right dorsal region to make an ulcer. The wound healing process was observed, and the re-epithelialization area was assessed stereoscopically.

### Re-epithelialization with keratinocyte transplantation

A conventional skin reconstitution method was performed with reference to previous studies.^13–15^ Four-week-old BALB/cAJcl-nu/nu female mice (Nippon Bio-Supp. Center) were used as recipient animals. Under inhalation anesthesia, a 6 mm circular piece of skin was excised from the right dorsal region. An autoclaved 1.0-cm-diameter silicone chamber, created using a 3D-printed template,^16^ was inserted into the skin hole. The rim of the silicone chamber was secured to the skin with four 5-0 nylon sutures. Ten million GFP-expressing keratinocytes were prepared in 150 μL of F medium and transferred into the chamber by making a small incision at the top. One week after transplantation, the upper half of the chamber was cut off. Two weeks after transplantation, the entire chamber was removed. The skin following re-epithelialization was evaluated 4 weeks after transplantation using a Stereoscope Zeiss AXIO Zoom V16 (Zeiss, Baden-Württemberg, Germany).

### Skin reconstruction through skin grafting

Non-pigmented skin in which the hair follicles were in the telogen stage^17^ was harvested from the dorsal part of GFP transgenic mice after hair clipping. The skin was thinned in two distinct thicknesses (5/1000 inch or 10/1000 inch) for the purpose of split-thickness skin grafting using a drum-type Padgett dermatome (Keisei Medical Industrial Co., Ltd., Tokyo, Japan). Under inhalation anesthesia, a longitudinal incision was made on the mid-dorsal back of 4-week-old BALB/cAJcl-nu/nu female mice and a subcutaneous pocket was created. Either of the two previously described types of donor skin or full-thickness donor skin was grafted onto the fascia and secured with 5-0 nylon sutures. The wound on the nude mouse was then closed using the surrounding skin. The covering skin was excised to expose the grafted area 5 days after transplantation. The sutures were removed 2 weeks after transplantation. Macroscopic and histological evaluation was conducted 5 weeks after transplantation. The skin tissue was harvested from the mid-dorsal region of the mouse, including both grafted and non-grafted areas. As autologous skin grafting, the same procedure, in which the mid-dorsal skin was harvested, thinned, and sutured at the same portion, was performed using single B6 mice in place of GFP transgenic mice and nude mice.

### Frozen sectioning

Tissues were fixed with 4% paraformaldehyde in PBS for 1 day and then incubated in 30% sucrose in PBS for 2 days. They were embedded in OCT compound and frozen with dry ice. Frozen samples were sectioned using a cryofilm [Cryofilm type 2C(9), 2.5 cm C-FP094; Section-Lab, Hiroshima, Japan] by a previously described method^18, 19^ at 200 μm intervals using a cryostat. Tissue sections were stained with hematoxylin and eosin (H&E) or mounted with 4′,6-diamidino-2-phenylindole (DAPI) Fluoromount-G (Thermo Fisher Scientific, Waltham, MA, USA). They were imaged with SLIDEVIEW VS200 Slide Scanner (Olympus, Tokyo, Japan).

### Statistical analysis

Numerical data are presented as mean ± standard deviation (SD). The significance of differences was examined using Welch’s t-test. Values of *P* < 0.05 were considered statistically significant.

## Results

### Evaluation of healed wound with or without keratinocyte transplantation

First, to develop a mouse model of acquired skin appendage dysfunction without any inherent abnormalities, wild-type mice (specifically B6 mice) were used. When skin ulcers were created by simple resection of circular skin pieces on the back of B6 mice, the wounds healed primarily through contraction of the surrounding skin within 2 weeks (Figure 1a). Given the limited area of re-epithelialization, the procedure was apparently insufficient to ensure a skin surface with a reduced number of skin appendages; therefore, we investigated the usefulness of transplanting cultured keratinocytes to the ulcer. To track the progression of wound area and engraftment of transplanted cells, allogeneic transplantation of primary keratinocytes isolated from the GFP transgenic mice (Figure 1b) to the wounds created on the back of immunodeficient nude mice was performed. This procedure allowed the external observation of areas of transplanted cell engraftments stereoscopically.

**Figure 1.**
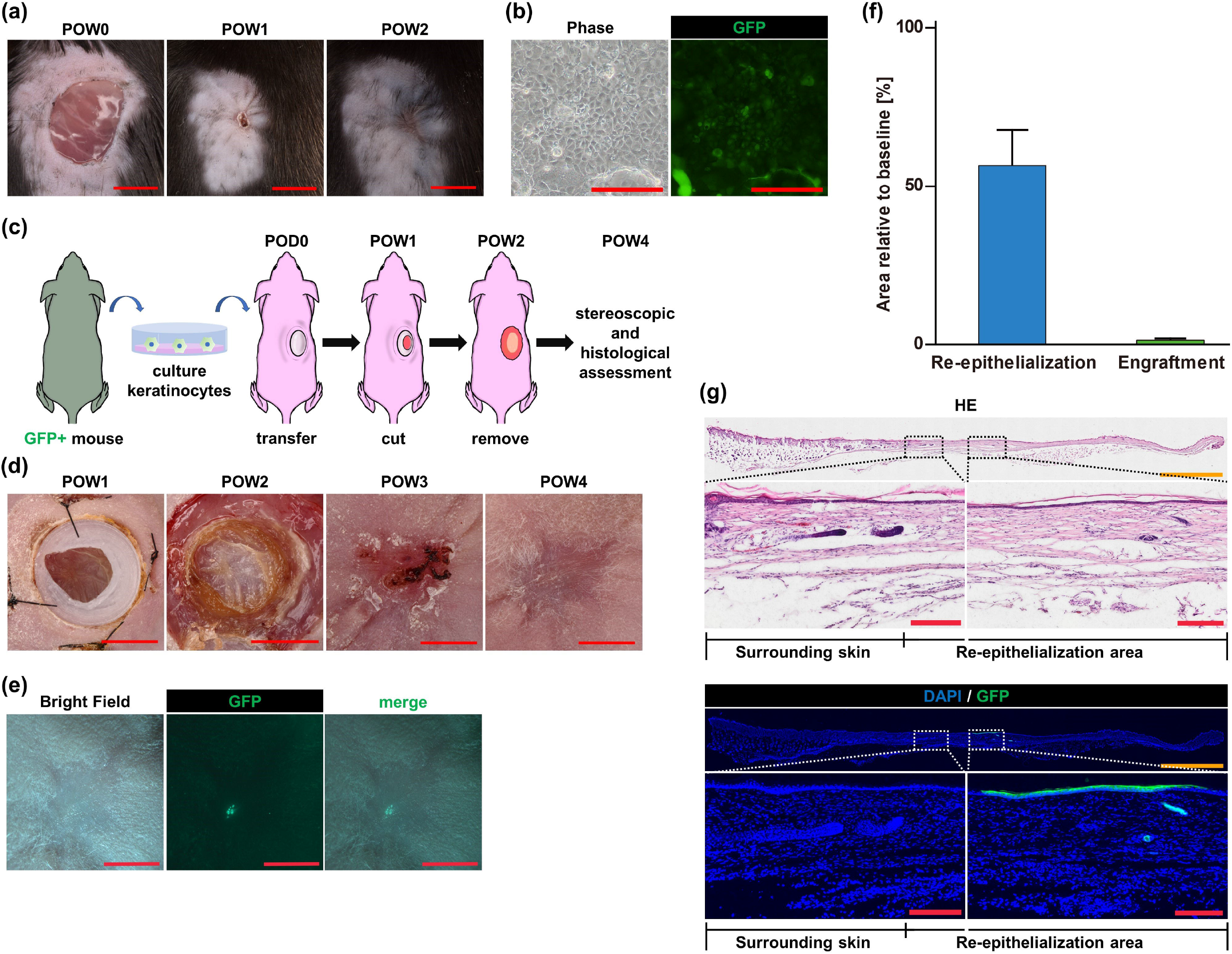
Evaluation of healed wound with or without keratinocyte transplantation. (a) Representative appearance of ulcers created by a 10 mm circular excision. By postoperative week (POW)2, complete re-epithelialization was achieved. Red scale bar, 5 mm. (b) Appearance of GFP-expressing keratinocytes on feeders at passage 2 from primary culture. Red scale bar, 200 μm. (c) Schematic image of the experimental design of re-epithelialization with GFP keratinocyte cell transplantation into a skin chamber on the back of nude mice. The silicone chamber was incised and opened at POW1 and removed at POW2. Stereoscopic analysis was conducted after re-epithelialization at POW4. (d) Representative appearance of ulcers following transplantation of GFP-expressing keratinocytes. By POW4, complete re-epithelialization was achieved. Red scale bar, 5 mm. (e) Representative appearance of GFP positivity after re-epithelialization. This indicated the presence of previously transplanted keratinocytes. Red scale bar, 5 mm. (f) The percentage of the areas of re-epithelialization and GFP-expressing keratinocyte engraftment. The area of a 10-mm-diameter silicone chamber was considered 100% (baseline) (mean ± SD; N=8). (g) Representative histological images of the area of re-epithelialization using hematoxylin and eosin (H&E) and DAPI staining. The presence of GFP expression in some epithelial cells indicated the successful engraftment of donor GFP-expressing keratinocytes, although the portion was limited. Orange scale bar, 2 mm; red scale bar, 200 μm.

A silicone chamber was attached to the ulcerated surface created by excision of the dorsal skin of a nude mouse, and keratinocytes were transplanted into the chamber. Two weeks after the cell transplantation, the chamber was removed and the site was monitored for another 2 weeks (Figure 1c). When the chamber was removed, epithelial-like tissue was observed covering the original ulcer area. The area was then covered with a scab, and by the fourth week, it was epithelialized in a concave shape compared to the surrounding area (Figure 1d). Stereoscopic observation revealed that only a small portion of the transplanted cells were located within the concavity (Figure 1e). The area of re-epithelialization, defined by depression relative to the surrounding skin, averaged 56.5 ± 11.3% when compared with the initial chamber area at the time of transplantation, while average GFP-positive area, equivalent to the area of engraftment of the transplanted cells, was only 1.31 ± 0.6% (Figure 1f). In addition to apparent lack of hair growth, histological evaluation revealed no evidence of skin appendages in the areas of re-epithelialization, regardless of whether the epithelium was composed of transplanted cells or recipient ones (Figure 1g).

In summary, keratinocyte transplantation could facilitate re-epithelialization to a certain extent without skin appendages, while the achieved area predominantly comprised recipient cells rather than transplanted ones.

### Evaluation of reconstructed skin after grafting of skin of varying thicknesses

To determine the usefulness of grafting of skin of different thicknesses to ensure a skin surface with a reduced number of appendages, skin portions obtained from the back of GFP transgenic mice were transplanted onto wounds created on the back of immunodeficient nude mice.

Mouse back skin dissected at the loose tissue layer was defined as a full-thickness skin graft. As a split-thickness skin graft, the skin was thinned in two distinct thicknesses (10/1000 inch or 5/1000 inch) using a drum-type Padgett dermatome (Figure 2a). Prior to transplantation, histological assessment was conducted to confirm the characteristics of the three types of donor skin, using H&E and DAPI staining. The full-thickness donor skin contained the epidermis, dermis, cutaneous muscle (panniculus carnosus),^20, 21^ underlying subcutaneous fat, and part of the muscle fascia. The 10/1000-inch donor skin included the epidermis, dermis, and subcutaneous tissue, with the cutaneous muscle, while the 5/1000-inch donor skin retained only the epidermis and dermis (Figure 2b).

**Figure 2.**
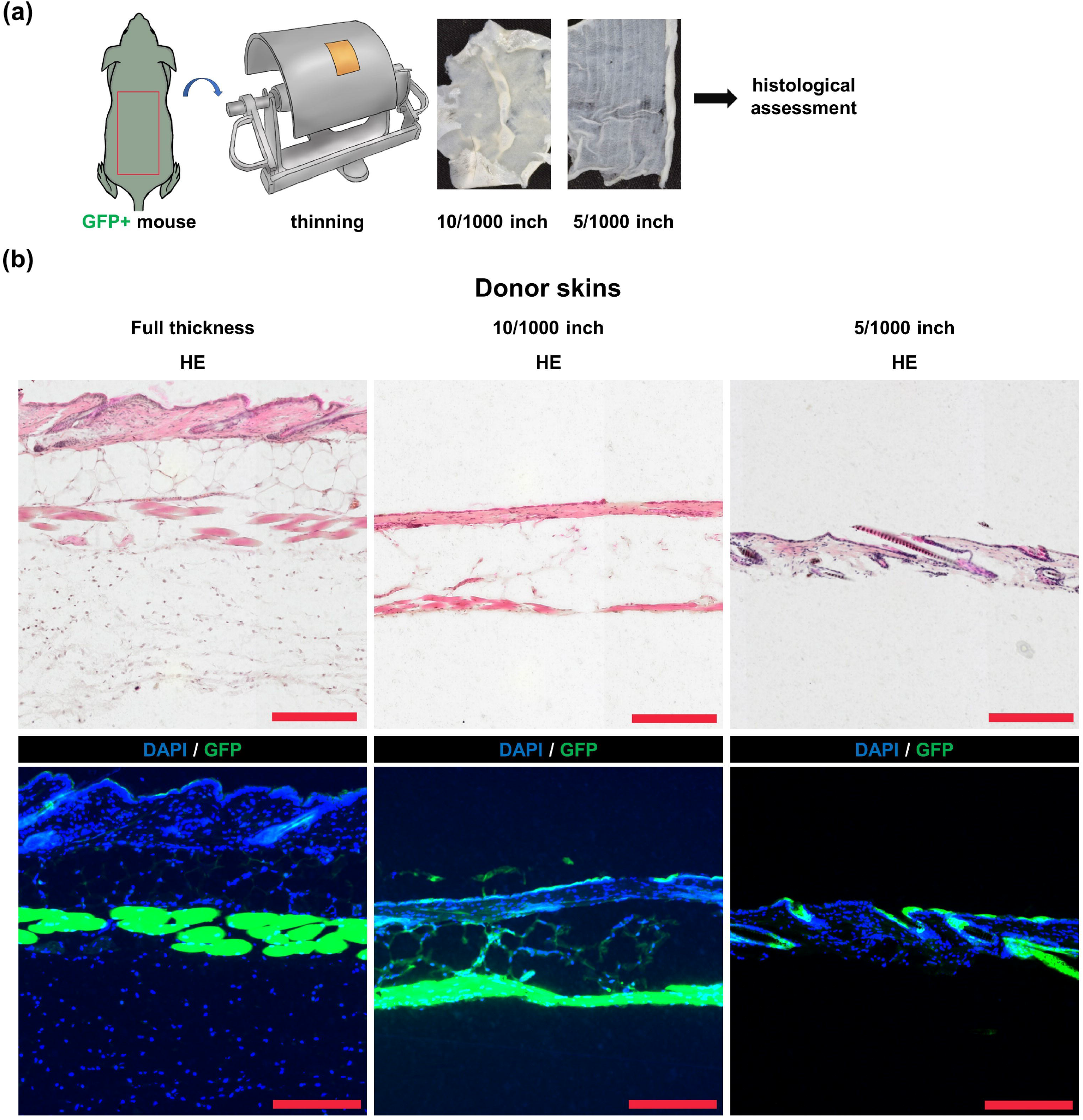
Evaluation of donor skin of varying thicknesses. (a) Schematic image of experimental design for making donor skin. Back skin of a GFP transgenic mouse dissected at the loose tissue layer was defined as a full-thickness skin graft. As a split-thickness skin graft, the skin was thinned in two distinct thicknesses (10/1000 inch or 5/1000 inch) using a drum-type Padgett dermatome. (b) Representative histological images of donor skin using hematoxylin and eosin (H&E) and DAPI staining. The full-thickness donor skin contained the epidermis, dermis, cutaneous muscle (panniculus carnosus), underlying subcutaneous fat, and some of the muscle fascia. The 10/1000-inch donor skin included the epidermis, dermis, and subcutaneous tissue, with the cutaneous muscle, while the 5/1000-inch donor skin retained only the epidermis and dermis. Red scale bar, 200 µm.

On transplantation, a longitudinal incision was made on the mid-dorsal back of the nude mice and a subcutaneous pocket was created. Processed graft was transplanted onto the fascia and secured with 5-0 nylon sutures. The wound on each nude mouse was then closed using the surrounding skin. The covering skin was excised 5 days after transplantation and the subsequent changes of each of the three types of grafted skin were evaluated in terms of appearance, engraftment area, and histology 5 weeks after transplantation (Figure 3a).

**Figure 3.**
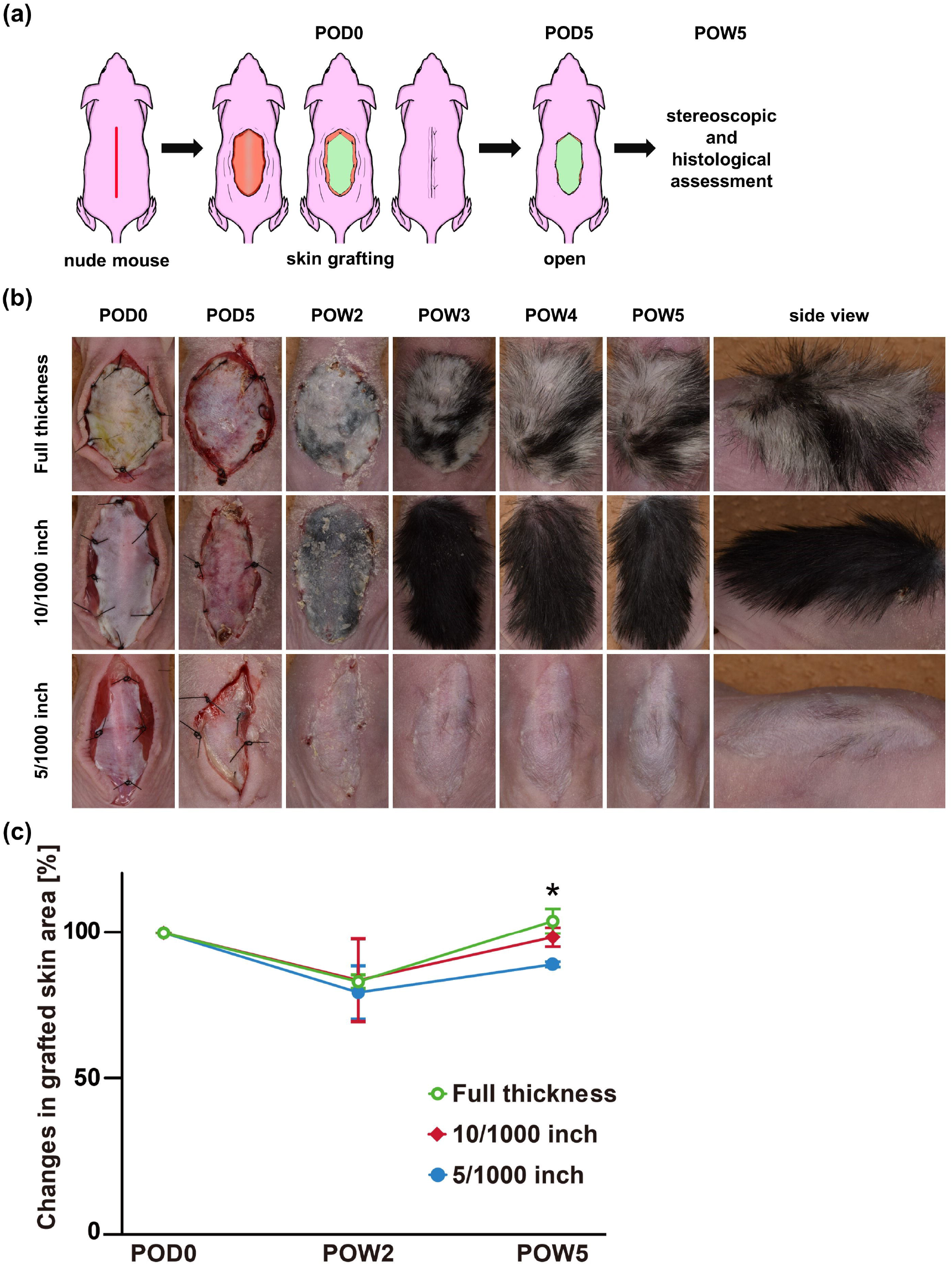
Macroscopic evaluation of reconstructed skin after grafting of GFP-expressing skin of varying thicknesses. (a) Schematic image of experimental design of skin reconstruction by GFP-expressing skin grafting on the back of nude mice. The 5/1000-inch and 10/1000-inch donor skin was prepared. A longitudinal incision was made on the mid-dorsal back of nude mice and a subcutaneous pocket was created. Either of the two previously described types of donor skin or full-thickness donor skin was grafted onto the fascia. The wound on the nude mouse was then closed using the surrounding skin. At postoperative day (POD)5, the covering skin was excised to expose the grafted area. Macroscopic and histological evaluations were conducted at POW5. (b) Representative appearance following grafting of GFP-expressing skin of each thickness. With the full-thickness graft, in general, the grafted skin turned whitish with black patches at 2 weeks, followed by growth of a moderate amount of white and black hair. With the 10/1000-inch graft, the grafted skin turned black at 2 weeks, followed by robust growth of black hair. In contrast, the 5/1000-inch graft consistently lacked pigmentation and showed little hair growth. (c) Changes in grafted skin area for each skin thickness. With the area at the time of skin grafting defined as 100%, average changes in size at POW2 and POW5 for each graft thickness are shown (mean ± SD; \**P* < 0.05 for full thickness vs. 5/1000 inch, and 10/1000 inch vs. 5/1000 inch; N=3 each).

The appearance of the grafted skin exhibited apparent differences across the three types of transplantations over the course of skin reconstruction (Figure 3b). With the full-thickness graft, in general, the grafted skin turned whitish with black patches at 2 weeks, followed by a moderate amount of growth of white and black hair. With the 10/1000-inch graft, the grafted skin turned black at 2 weeks, followed by robust growth of black hair. Interestingly, the pigmentation occurred at precisely 2 weeks in all mice. In contrast, the 5/1000-inch graft consistently lacked pigmentation and showed little hair growth. It is important to note that the distribution of hair was not uniform. Some areas demonstrated a complete absence of hair, while others exhibited some hair, albeit at a low density.

To examine the degree of grafted skin contraction, the grafted area was measured and compared at 2 and 5 weeks after transplantation (Figure 3c). With the area at the time of skin grafting being defined as 100%, the areas of the full-thickness, 10/1000-inch, and 5/1000-inch grafts were reduced down to 83.8 ± 2.26%, 84.3 ± 13.7%, and 80.2 ± 8.86% 2 weeks after transplantation, and then increased to 104.0 ± 4.1%, 99.2 ± 3.2%, and 89.5 ± 0.9% 5 weeks after transplantation, respectively.

To determine whether the grafted skin had been successfully transplanted or merely exhibited re-epithelialization due to recipient tissue, and particularly to assess the decreased number of skin appendages, histological assessments of reconstructed skin at 5 weeks were performed (Figure 4a). While only DAPI staining was observed in the non-grafted areas, GFP was also visible in the grafted areas, clearly delineating their boundaries. The hair follicles were moderately visible in the section of full-thickness skin grafts. A large number of hair follicles were visible in the 10/1000-inch section, while there was slight thickening of the fibrous tissue with minimal visible hair follicles in the 5/1000-inch section. For comparison, the number of hair follicles was determined for each skin thickness (Figure 4b). For each thickness, hair follicles were first counted in five sections, each 5 mm in length, and the average was calculated. This process was repeated for three mice, and the overall average number of hair follicles was determined. Additionally, the number of hair follicles in the full-thickness donor skin, which served as a control, was similarly determined. Full-thickness sections had slightly fewer hair follicles, with a mean of 42.4 ± 12.9, compared with the 10/1000-inch sections and the control sections, which contained 63.2 ± 9.46 and 69.7 ± 15.8, respectively, although these differences were not statistically significant. In contrast, 5/1000-inch sections exhibited markedly fewer hair follicles, with a mean of 3.73 ± 4.46 (*P* < 0.05). It should be noted that the distribution of hair follicles in 5/1000-inch sections was not uniform, as observed macroscopically. The histological areas exhibiting absolutely no hair follicles corresponded to the macroscopically hairless areas, and the histological areas with some density of hair follicles overlapped with the macroscopically hair-growing areas.

**Figure 4.**
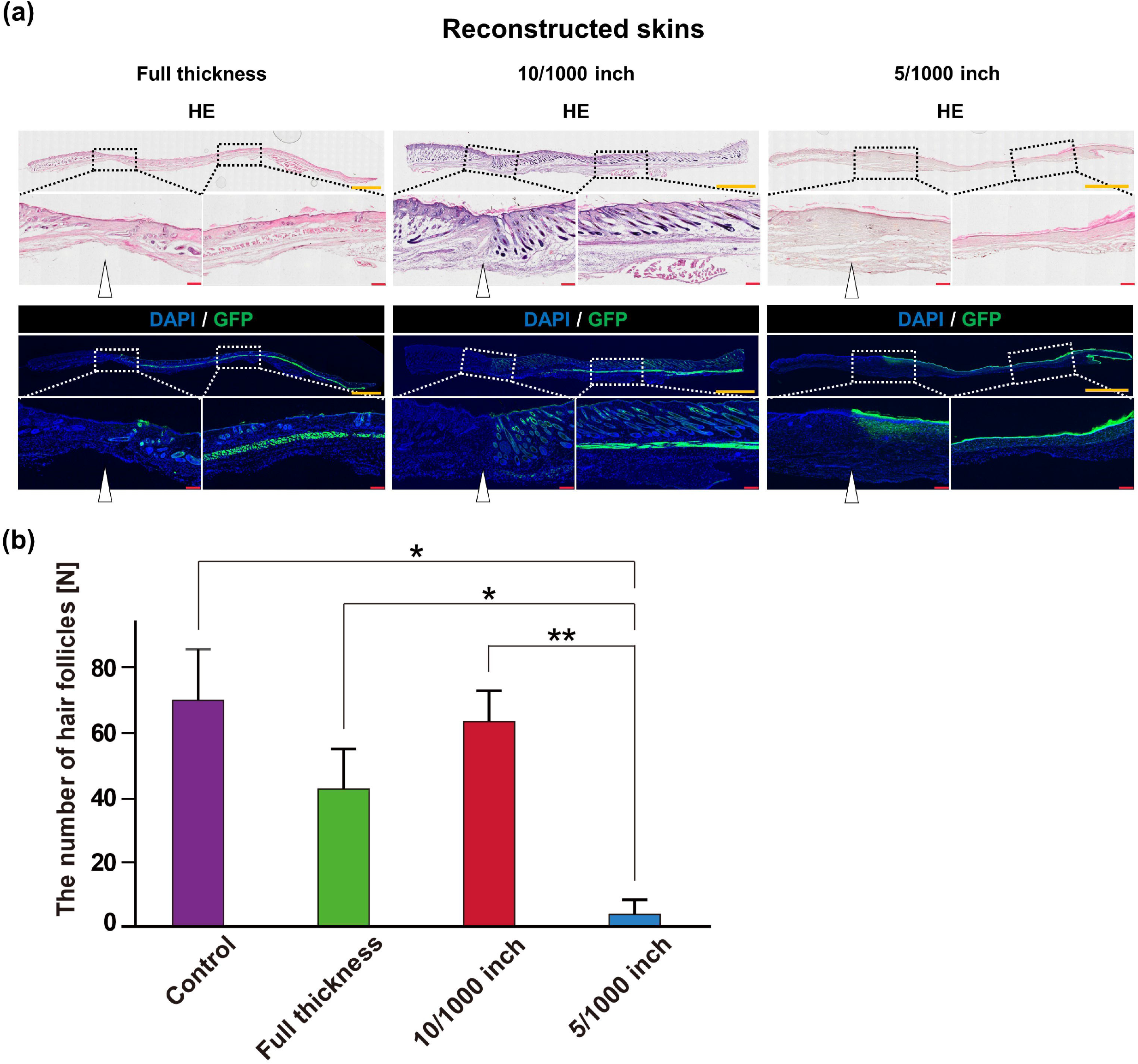
Histological evaluation of reconstructed skin after grafting of GFP-expressing skin of varying thicknesses. (a) Representative histological images of reconstructed skin using hematoxylin and eosin (H&E) and DAPI staining. In the section of full-thickness skin grafts, the hair follicles were moderately visible. A large number of hair follicles were visible in the 10/1000-inch section, while there was slight thickening of the fibrous tissue with minimal visible hair follicles in the 5/1000-inch section. Orange scale bar, 2 mm; red scale bar, 200 µm; white arrow, the boundary between surrounding and grafted skin. (b) The hair follicle count for each skin thickness. For each thickness, hair follicles were first counted in five sections, each 5 mm in length, and the average was calculated. This process was repeated for three mice, and the overall averages were determined. Additionally, the number of hair follicles in full-thickness donor skin, serving as a control, was similarly quantified (mean ± SD; \**P* < 0.05, \*\**P* < 0.01; N=3 each).

Finally, the usefulness of 5/1000-inch skin grafting for generating skin surfaces with a reduced number of skin appendages was tested in orthotopic autologous transplantation (Figure 5a). The autologous 5/1000-inch grafted skin consistently lacked pigmentation and showed little hair growth at 5 weeks, comparable to the outcomes observed in allogeneic skin grafting (Figure 5b). Histological assessment of reconstructed skin at 5 weeks revealed the presence of minimal skin appendages with a slight increase in dermal thickness, which was also consistent with the characteristics of allogeneic skin grafting (Figure 5c).

**Figure 5.**
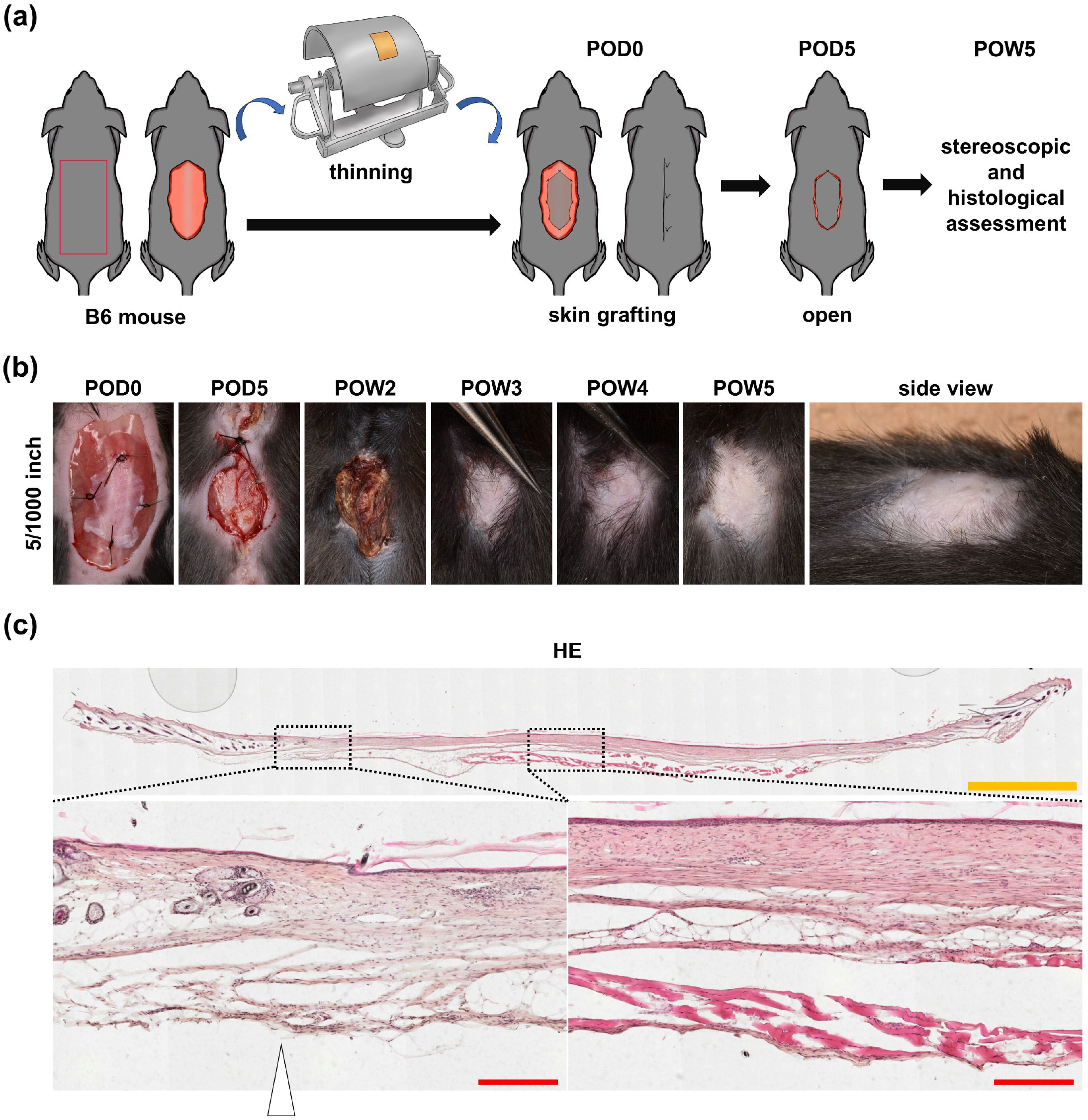
Evaluation of reconstructed skin after autologous 5/1000-inch skin grafting. (a) Schematic image of experimental design of skin reconstruction by autologous 5/1000-inch skin grafting. The mid-dorsal skin of B6 mice was harvested, thinned to a thickness of 5/1000 inch, and sutured at the same portion. On postoperative day (POD)5, the covering skin was excised to expose the grafted area. Macroscopic and histological evaluations were conducted at POW5. (b) Representative appearance following autologous 5/1000-inch skin grafting. The grafted skin consistently lacked pigmentation and showed little hair growth at POW5. (c) Representative histological images of reconstructed skin at POW5 using hematoxylin and eosin (H&E) staining. Minimal skin appendages with a slight increase in dermal thickness were observed. Orange scale bar, 2 mm; red scale bar, 200 µm; white arrow, the boundary between surrounding and grafted skin.

## Discussion

In this study, we assessed methodologies for developing a mouse model with a reduced number of skin appendages. The keratinocyte transplantation enabled a certain skin area without skin appendages to be achieved, which could not be obtained by simple skin wounding. One important aspect of this method is that no regeneration of skin appendages was confirmed, at least within the period that we examined. If a relatively large area of skin completely lacking skin appendages is needed, keratinocyte transplantation could be a good option with the assumption that the regeneration of skin appendages over a longer period should be confirmed according to the needs of the planned research. Additionally, it is noteworthy that most of the epithelialized areas obtained were composed of epithelial cells derived from the surrounding recipient epithelium, rather than from the transplanted cells. The method is not recommended when epithelium composed of transplanted cells is required (e.g., human cell xenograft to evaluate the regenerative capacity of human-derived cells), although exploring various cell transplantation conditions, such as optimization of the number of transplanted keratinocytes, altering the transplantation medium, and co-transplantation with mesenchymal cells, may enhance cell engraftment rates.

Meanwhile, skin reconstruction by skin grafting demonstrated stability and the feasibility of extensive transplantation, enabling the production of skin with minimal skin appendages macroscopically and histologically when adjusted to 5/1000-inch thickness. The 5/1000-inch grafted skin exhibited greater contraction than those of other thicknesses, but maintained approximately 90% of the original grafted area, ensuring an area sufficiently large for evaluating the regeneration of skin appendages. The 5/1000-inch donor skin has a thin part of the dermis shaved off and does not include the dermal papilla; therefore, less hair growth is observed. When harvesting donor skin, it is essential to consider the phase of the hair cycle, which is categorized as anagen, catagen, and telogen. The localization of hair follicles between cutaneous muscle and dermis as well as the thickness of subcutaneous fat and dermis changes over the course of the hair cycle. The skin turns black in anagen and pink in telogen.^17^ The telogen phase is divided into two distinct categories: refractory telogen and competent telogen. During the refractory telogen phase, the presence of bone morphogenetic proteins (BMPs) within the surrounding skin inhibits activation of the hair follicles, preventing the onset of hair regeneration.^22^ Conversely, the absence of BMPs during the competent telogen phase allows initiation of the anagen phase when Wingless-related MMTV integration site (WNT) is activated.^22^ A previous study showed that skin grafting, even in refractory telogen, can achieve hair growth.^23^ Thus, donor skin in telogen, regardless of these two phases, has the potential for hair regeneration and was harvested and evaluated in this study. One interesting finding is that all grafted skin, especially in the 10/1000-inch section with a high concentration of hair follicles, exhibited blackening and the emergence of hair 2 weeks after transplantation regardless of the timing of the telogen phase of donor skin. This may indicate that the hair follicles across the grafted skin underwent simultaneous anagen re-entry and multiple hair cycle areas reset into a single area, which is recognized as trauma-caused anagen.^24^ This phenomenon can result from external factors such as the expression of S100 proteins, Map3k5-dependent infiltration, and activation of peri-wound macrophages.^25, 26^ The anagen stage typically lasts 1–3 weeks. Thus, if hairs do not emerge in some areas 3 weeks after transplantation, it is highly likely that the hair follicles are absent in the area. This assumption is consistent with the results observed in the 5/1000-inch grafted skin 5 weeks after transplantation, in which the histological areas devoid of hair follicles corresponded to the macroscopically hairless areas. Although achieving complete hairlessness of the entire skin is challenging, successful creation of completely hairless skin in some areas is relatively easy, so a mouse model with skin grafting is considered to have high utility as an experimental model. Furthermore, it was considered possible to apply this method to xenografts using human skin tissue, although some adjustment of thickness could be necessary. Our recent study demonstrated that transplantation of a specific type of epithelial cell (introduced with LEF1) and two types of mesenchymal cells (introduced with FOXD1 and PRDM1, or with LEF1 and SHH), which were all reprogrammed from subcutaneous mesenchymal cells of adult mice to mimic developing skin cells, to a skin ulcer could lead to the formation of skin appendage-like structures.^12, 27, 28^ This mouse model is intended to be used to assess the efficacy of introducing the genes involved to achieve hair follicle regeneration.

In conclusion, to obtain skin with minimal skin appendages, a mouse model with keratinocyte transplantation is advantageous when it is acceptable for the model to be created through allogeneic transplantation and re-epithelialization is achieved by the surrounding epidermis rather than transplanted cells. Meanwhile, a mouse model with skin grafting at a thickness of 5/1000 inch is widely applicable due to the fact that the model can be created through autologous transplantation and xenotransplantation, as well as allogeneic transplantation. These two mouse models of acquired skin appendage dysfunction have the potential to serve as valuable research tools for the development of novel treatments aimed at regenerating skin appendages.

## Acknowledgments

We thank Edanz (https://jp.edanz.com/ac) for editing a draft of this manuscript. This work was supported by JSPS KAKENHI grant number JS22H03247 (to M.K. and M.O.), JP20K20609 (to M.K. and M.O.), and AMED under grant numbers JP20bm070403 (to M.K.) and JP21zf0127002 (to M.K.).

## Ethics statement

All animal experiments were approved by the Animal Research Committee of the University of Tokyo.

## References

1. Millar SE. Molecular mechanisms regulating hair follicle development. J Invest Dermatol. 2002 Feb;118(2):216–25.

2. Sennett R, Wang Z, Rezza A, Grisanti L, Roitershtein N, Sicchio C, et al. An Integrated Transcriptome Atlas of Embryonic Hair Follicle Progenitors, Their Niche, and the Developing Skin. Dev Cell. 2015 Sep 14;34(5):577–91.

3. Gilhar A, Ullmann Y, Berkutzki T, Assy B, Kalish RS. Autoimmune hair loss (alopecia areata) transferred by T lymphocytes to human scalp explants on SCID mice. J Clin Invest. 1998 Jan 1;101(1):62–7.

4. Yazdabadi A, Magee J, Harrison S, Sinclair R. The Ludwig pattern of androgenetic alopecia is due to a hierarchy of androgen sensitivity within follicular units that leads to selective miniaturization and a reduction in the number of terminal hairs per follicular unit. Br J Dermatol. 2008 Dec;159(6):1300–2.

5. Cash TF. The psychology of hair loss and its implications for patient care. Clin Dermatol. 2001;19(2):161–6.

6. Zheng Y, Du X, Wang W, Boucher M, Parimoo S, Stenn K. Organogenesis from dissociated cells: generation of mature cycling hair follicles from skin-derived cells. J Invest Dermatol. 2005 May;124(5):867–76.

7. Zhao Q, Li N, Zhang H, Lei X, Cao Y, Xia G, et al. Chemically induced transformation of human dermal fibroblasts to hair-inducing dermal papilla-like cells. Cell Prolif. 2019 Sep;52(5):e12652.

8. Mäkelä OJM, Mikkola ML. Mesenchyme governs hair follicle induction. Development. 2023 Nov 15;150(22).

9. Benavides F, Oberyszyn TM, VanBuskirk AM, Reeve VE, Kusewitt DF. The hairless mouse in skin research. J Dermatol Sci. 2009 Jan;53(1):10–8.

10. Sander AL, Jakob H, Henrich D, Powerski M, Witt H, Dimmeler S, et al. Systemic transplantation of progenitor cells accelerates wound epithelialization and neovascularization in the hairless mouse ear wound model. J Surg Res. 2011 Jan;165(1):165–70.

11. Sommer K, Jakob H, Kisch T, Henrich D, Marzi I, Frank J, et al. Local application reduces number of needed EPC for beneficial effects on wound healing compared to systemic treatment in mice. Eur J Trauma Emerg Surg. 2022 Jun;48(3):1613–24.

12. Kurita M, Araoka T, Hishida T, O’Keefe DD, Takahashi Y, Sakamoto A, et al. In vivo reprogramming of wound-resident cells generates skin epithelial tissue. Nature. 2018 Sep;561(7722):243–7.

13. Yuspa SH, Morgan DL, Walker RJ, Bates RR. The growth of fetal mouse skin in cell culture and transplantation to F1 mice. J Invest Dermatol. 1970 Dec;55(6):379–89.

14. Worst PK, Mackenzie IC, Fusenig NE. Reformation of organized epidermal structure by transplantation of suspensions and cultures of epidermal and dermal cells. Cell Tissue Res. 1982;225(1):65–77.

15. Lichti U, Anders J, Yuspa SH. Isolation and short-term culture of primary keratinocytes, hair follicle populations and dermal cells from newborn mice and keratinocytes from adult mice for in vitro analysis and for grafting to immunodeficient mice. Nat Protoc. 2008;3(5):799–810.

16. Du Z, Shen Q, Mito D, Kato M, Okazaki M, Kurita M. Optimized 3D-printed template design for production of silicone skin chambers. J Dermatol Sci. 2022 Jan 1;105(1):55–7.

17. Müller-Röver S, Handjiski B, van der Veen C, Eichmüller S, Foitzik K, McKay IA, et al. A comprehensive guide for the accurate classification of murine hair follicles in distinct hair cycle stages. J Invest Dermatol. 2001 Jul;117(1):3–15.

18. Kawamoto T, Kawamoto K. Preparation of thin frozen sections from nonfixed and undecalcified hard tissues using Kawamot’s film method (2012). Methods Mol Biol. 2014;1130:149–64.

19. Shen Q, Suga S, Moriwaki Y, Du Z, Aizawa E, Okazaki M, et al. Optimization of an adeno-associated viral vector for keratinocytes in vitro and in vivo. Available from: 10.1101/2024.04.15.589645

20. Bahri OA, Naldaiz-Gastesi N, Kennedy DC, Wheatley AM, Izeta A, McCullagh KJA. The panniculus carnosus muscle: A novel model of striated muscle regeneration that exhibits sex differences in the mdx mouse. Sci Rep. 2019 Nov 4;9(1):15964.

21. Imazato H, Takahashi N, Hirakawa Y, Yamaguchi Y, Hiyoshi M, Tajima T, et al. Three-dimensional fine structures in deep fascia revealed by combined use of cryo-fixed histochemistry and low-vacuum scanning microscopy. Sci Rep. 2023 Apr 18;13(1):6352.

22. Plikus M V., Baker RE, Chen CC, Fare C, de la Cruz D, Andl T, et al. Self-Organizing and Stochastic Behaviors During the Regeneration of Hair Stem Cells. Science (1979). 2011 Apr 29;332(6029):586–9.

23. Plikus M V., Mayer JA, de la Cruz D, Baker RE, Maini PK, Maxson R, et al. Cyclic dermal BMP signalling regulates stem cell activation during hair regeneration. Nature. 2008 Jan;451(7176):340–4.

24. Plikus M V, Chuong CM. Complex hair cycle domain patterns and regenerative hair waves in living rodents. J Invest Dermatol. 2008 May;128(5):1071–80.

25. Ito M, Kizawa K. Expression of calcium-binding S100 proteins A4 and A6 in regions of the epithelial sac associated with the onset of hair follicle regeneration. J Invest Dermatol. 2001 Jun;116(6):956–63.

26. Osaka N, Takahashi T, Murakami S, Matsuzawa A, Noguchi T, Fujiwara T, et al. ASK1-dependent recruitment and activation of macrophages induce hair growth in skin wounds. J Cell Biol. 2007 Mar 26;176(7):903–9.

27. Kurita M, Izpisua Belmonte JC, Suzuki K, Okazaki M. Development of de novo epithelialization method for treatment of cutaneous ulcers. J Dermatol Sci. 2019 Jul 1;95(1):8–12.

28. Moriwaki Y, Qi S, Okada H, Zening D, Suga S, Kato M, et al. In vivo reprogramming of wound-resident cells generates skin with hair. https://www.biorxiv.org/content/10.1101/2023.03.05.531138v1

